# Analyzing the Molecular Mechanism of Eucalyptol, Limonene and Pinene Enteric Capsules (QIENUO) in the Treatment of Pulmonary cystic fibrosis with Network Pharmacology and Verifying Molecular Docking

**DOI:** 10.1101/2024.09.30.615978

**Authors:** Tiantaixi Tu, Xinjie Zhu, Congyin Wang, Wei Chen, Yihu Zheng

**Affiliations:** Translational Medicine Laboratory, the First Affiliated Hospital of Wenzhou Medical University, China; Renji College, Wenzhou Medical University, China; The First School of Medicine, Wenzhou Medical University, China; Department of Surgery, the First Affiliated Hospital of Wenzhou Medical University, China

**Keywords:** Network Pharmacology, Molecular Docking, Pulmonary cystic fibrosis, Eucalyptol, Limonene and Pinene Enteric Capsules

## Abstract

**Objective:** To explore the mechanism of Eucalyptol, Limonene and Pinene Enteric Capsules (QIENUO) in the treatment of pulmonary cystic fibrosis (CPF), analyze the common targets of QIENUO and CPF, and verify the molecular docking of core proteins and small molecules.

**Methods:** The main active compounds and their corresponding targets were obtained from PubChem, SwissTargetprediction, GeneCards, PharmMapper and TCMSP databases. Targets related to CPF were screened from GeneCards, OMIM, DisGeNET and TTD databases. The common “QIENUO-CPF” targets were analyzed by gene ontology (GO) and Kyoto Encyclopedia of Genes and Genomes (KEGG) through the website of Weishengxin. Protein-protein interaction (PPI) network and compound-target-pathway network were constructed by Cytoscape, and the network parameters were systematically analyzed. The interaction between core protein and monomer components was evaluated and verified by molecular docking method.

**Results:** 228 active compounds target and 1354 CPF-related targets were screened out, and 92 common targets were analyzed by GO and KEGG. The results showed that the therapeutic effect of QIENUO on CPF was mainly through AMPK signaling pathway, cGMP-PKG signaling pathway and TGF-β signaling pathway. The results of molecular docking show that the binding energy of 9 of 15 pairs of ligand-receptor pairs is lower than-6 kjmol-1.

**Conclusion:** QIENUO exhibits huge potential as a therapeutic agent for the treatment of pulmonary cystic fibrosis. The specific molecular mechanism and effective active components of QIENUO treat CPF were studied and demonstrated, which provided theoretical basis for better clinical application of QIENUO.

## 1. Introduction

Cystic fibrosis is an autosomal recessive genetic disease, which is mainly caused by the pathogenic variation of cystic fibrosis transmembrane conductance regulator gene, which leads to the increase of mucus viscosity and the difficulty of mucus clearance. The main clinical manifestation of patients with pulmonary cystic fibrosis (CPF) is pulmonary obstructive symptoms^[1]^, it is characterized by progressive bronchiectasis and impaired lung function, accompanied by severe airflow obstruction. Respiratory failure is the main reason for shortening the life expectancy of patients with CPF. The pathogenesis of CPF is the result of the interaction of many internal and external factors, including genotype, mucus composition, abnormal movement of cilia, chronic inflammation and chronic airway infection. The risk of pulmonary complications in patients with CPF increases significantly with the progress of pulmonary diseases^[2]^. For example, once a patient with CPF is infected with Pseudomonas aeruginosa, it will lead to accelerated lung function impairment and seriously affect the prognosis and quality of life. Pseudomonas aeruginosa showed significant resistance to antibiotics^[3-8]^. Inhaled tobramycin is an important treatment^[9]^. Phlegm-resolving drugs are important auxiliary drugs for the treatment of complications of CPF. Myrtle oil^[10,11]^ and Eucalyptol, Limonene and Pinene Enteric Capsules (QIENUO) are the two main drugs for transforming phlegm, refining from plants. Standardized Myrtle is an essential oil mainly containing cineole, limonene and α-pinene. Studies have proved that standardized Myrtle can alleviate acute lung injury induced by lipopolysaccharide in mice, and it is mainly used to treat sinusitis, bronchitis and chronic obstructive pulmonary disease in clinic^[12]^. QIENUO are used as expectorants in clinic and widely used to treat various respiratory diseases, including chronic bronchitis^[13]^. It has been reported before that they may play a therapeutic role by inhibiting TLR4 signal transduction^[14]^.

Network pharmacology was first put forward by Hopkins, which is the target of mapping multi-pharmacological network to human disease-gene network and revealing drug action^[15]^. In our study, the active compounds and corresponding targets in QIENUO were screened by introducing databases of TCMSP, PubChem, GeneCards, PharmMapper and UniProt. Combined with OMIM, GeneCards, DisGENT and TTD databases, the relevant targets of CPF were screened. Through PPI database, the interaction of QIENUO-CPF common target was found. Enrichment analysis was used to evaluate the distribution of common targets of QIENUO -CPF in diseases, cellular components (CC), biological processes (BP), molecular functions (MF) and signal pathways. Finally, the network model of “drug-compound-conformation-target-gene” in the treatment of CPF with QIENUO was put forward. The purpose of this study is to explore how QIENUO can treat CPF through network pharmacology, and to verify it by molecular docking^[16-18]^. These findings provide a new perspective for treating CPF with traditional medicinal plants, and provide theoretical guidance for further experimental research and clinical application.

## 2. Materials and methods

### 2.1. Screening of active components and their action targets in QIENUO

The active components and their interacting proteins of QIENUO were identified by using the database of traditional Chinese medicine system pharmacology and analysis platform (TCMSP: http://tcmspw.com/tcmsp.php), and the screening criteria were established based on two basic pharmacokinetic indexes: oral bioavailability (OB) and drug similarity (DL), and the thresholds were set as OB≥30% and DL≥0.18. A free chemical structure database (PubChem: pubchem.ncbi.nlm.nih.gov/) maintained by the National Biotechnology Information Center (NCBI) under the National Institutes of Health (NIH), Target Prediction Database Swiss Target Prediction (http://www.swisstargetprediction.ch/) Comprehensive database of human genes (GeneCards: www.genecards.org/), The pharmacophore matching and potential target identification platform (PharmMapper: www.lilab-ecust.cn/pharmmapper/) developed and maintained by East China University of Science and Technology uses the universal protein resource database (UniProt: https://www.uniprot.org/) to convert the target protein names into official gene symbols.

### 2.2. Collection of targets related to CPF

Aim to determine the potential target of CPF, information was searched in four online databases: online mendelian inheritance in man (OMIM: https://omim.org/). GeneCards: the human gene database (GeneCards: https://www.genecards.org/ and therapeutic target database (http://db.idrblab.net/ttd/). DisGENT (DisGENT: https://disgenet.com/). These databases contain information about human genes and genetic diseases, as well as potential therapeutic targets of diseases. By integrating all targets and deleting duplicate data, all relevant targets can be obtained.

### 2.3. Common target overlap between active compound and CPF related target

By identifying the common target shared by the target related to CPF and the predicted QIENUO target, the results were displayed in venn diagram by using the VENNY2.1 website (https://bioinfogp.cnb.csic.es/tools/Venny/index.html).

### 2.4. Building a visual PPI network

Known to be both drug targets and disease genes targets are imported into string (http://string-db.org) database, which is a tool to find interacting genes and protein. The database is set to “Homosapiens” species, and the confidence of PPI is set to 0.7. Use Cytoscape software (version 3.8.2) to create a network of potential key targets. Then use CytoHubba plug-in to screen the core targets in the network. The core goal color is determined according to the Degree Centrality (DC) of the basic topological parameter, and the darker the color in the diagram, the greater the DC. Molecular Complexity Detection (MCODE) algorithm plug-in is used to detect and analyze the module structure in biological networks.

### 2.5. Enrichment analysis of DO, GO and KEGG

The enrichment analysis of disease ontology (DO), gene ontology (GO) and Kyoto Encyclopedia of Genes and Genomes (KEGG) was carried out by using DAVID database (David: https://david.ncifcrf.gov/) to evaluate the common “drug-disease” targets. In the enrichment analysis, the visual analysis was carried out through the website of weishengxin website (https://www.bioinformatics.com.cn/login/.

### 2.6. Construction of Topological Network

In order to clarify the relationship between the active compounds of QIENUO and the “drug-disease” target, a “drug-active compound-target” network was established by using Cytoscape version 3.8.2, and the topology analysis and visualization were carried out. The “drug-compound-target-pathway” network was also created and analyzed by Cytoscape, which intuitively showed the complex interactions among drugs, compounds, targets and pathways. Show the top five core proteins by R package.

### 2.7. Molecular docking verification

Obtain the core ligand structure file (mol format) from the TCMSP database (http:/http://tcmspw.com/tcmsp.php). The three-dimensional crystal structure of the target was obtained from the RCSB protein database (RCSB database: https://www.pdb.org/), and then was introduced into PyMOL for ligand separation. Both ligands and receptors are imported into AutoDockTools 1.5.6 (https://ccsb.scripts.edu/mgltools/downloads/) for dehydration, hydrogenation and charge calculation, and then stored in pdbqt format as a preparation step before molecular docking. The grid box for docking is created in AutoDockTools, including the whole target protein, and the parameters are saved in txt format. Finally, the molecular docking analysis was carried out by using AutoDockVina1.1.2 version 1.1.2 (https://vina.scripps.edu), and the free binding energy was visualized as a heat map by using the website of Microsense (Microsense: https://www.bioinformatics.com.cn/login/. The visualization of molecular docking and its interaction is displayed by PyMOL.

## 3. Results

### 3.1. Intersecting targets of QIENUO and CPF

A total of 228 potential gene targets related to QIENUO were found from three components of QIENUO collected from PubChem, SwissTargetprediction, GeneCards, PharmMapper and TCMSP databases. By integrating the targets of CPF collected from GeneCards, OMIM, DisGeNET and TTD, and removing the repeated targets, 1354 disease gene targets related to CPF were finally determined. In venn diagram, 92 common targets were identified as common targets with the active compound target of QIENUO and the target of CPF (Fig. 2A). These common targets may be important targets for the treatment of CPF with QIENUO. A “drug-compound-target” network consisting of three components and 92 targets was constructed by Cytoscape. Lines in the network represent the relationship between compounds and targets (Fig. 2B). The three components of QIENUO and their corresponding genes are shown here(Fig. 2C).

**Fig. 1.**
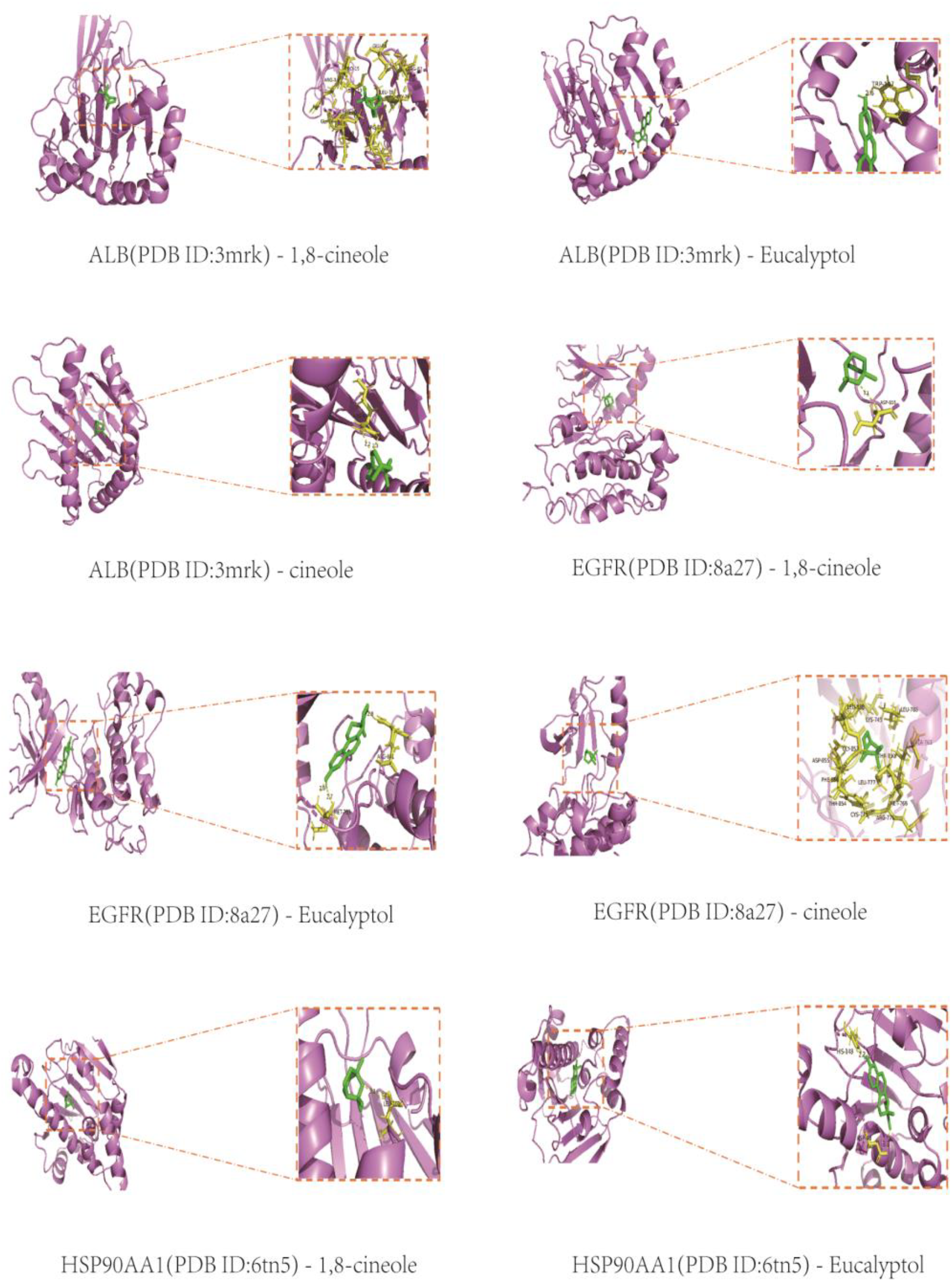
The ideas of the article and the main methods used are shown in the flowchart.

**Fig. 2.**
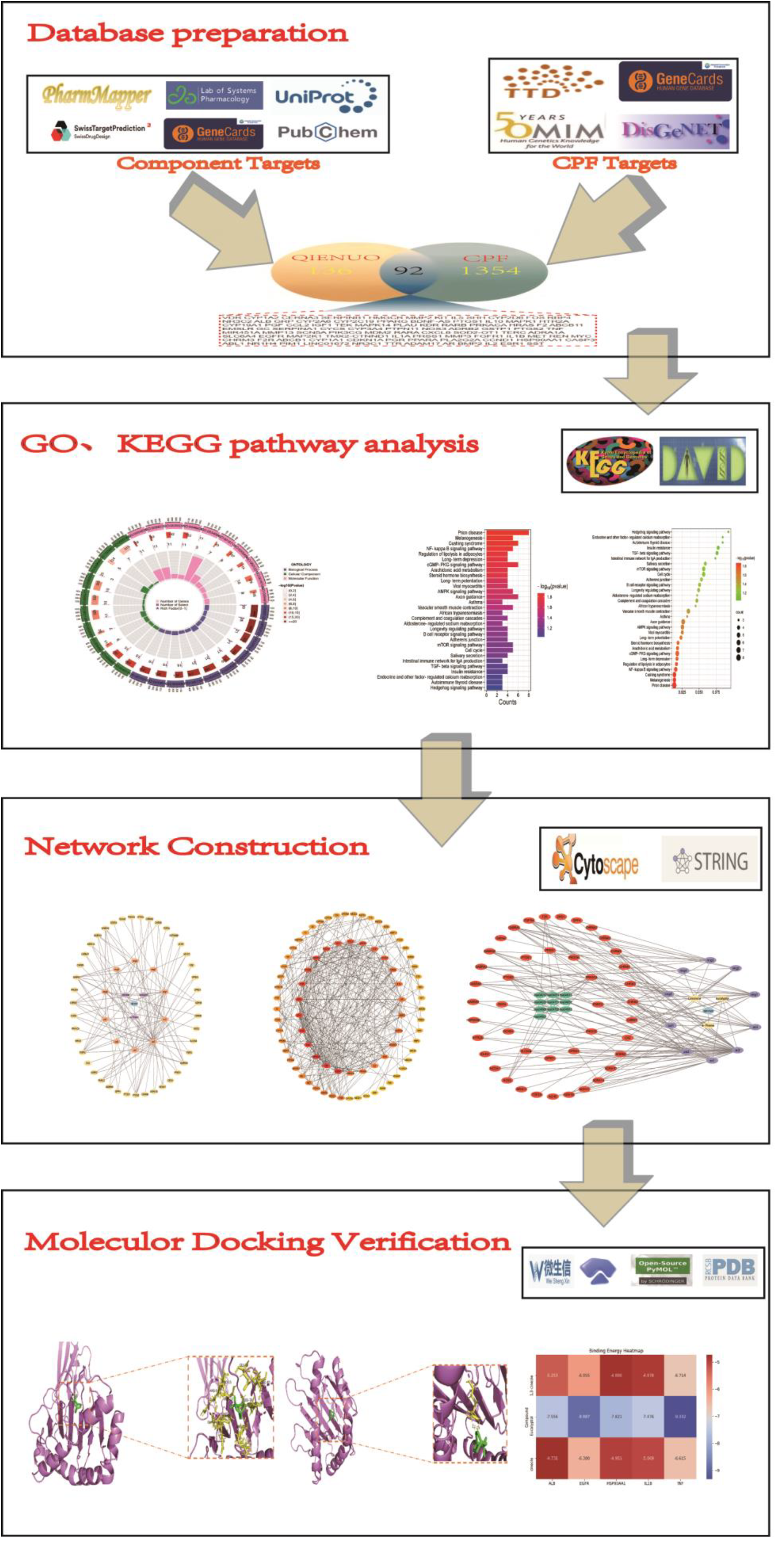
Screening of QIENUO active compounds and targets in CPF, Construction of drug-compound-target network. (A) Venn diagram of 268 common targets between active compound targets of Eucalyptol, Limonene and Pinene Enteric Capsules and CPF disease targets.(B) drug-compound-target network.

### 3.2. Analysis and Construction of Common Target PPI Network

a PPI network (Fig. 3A) based on 92 common targets was constructed by using STRING and Cytoscape. DC value is the most direct indicator of node centrality. The color of the node in the figure has changed from light to dark, indicating that the DC value of the node increases from low to high. In addition, the MCODE plug-in in Cytoscape is clustered, a highly connected subnet is constructed, and the targets are divided into five categories (Fig. 3B). Seventeen core objectives were determined by using the screening criteria based on topology analysis, and the threshold was ≥2 times DC (Fig. 3C). The top 17 core objectives are displayed in the form of a histogram (Fig. 3D).

**Fig. 3.**
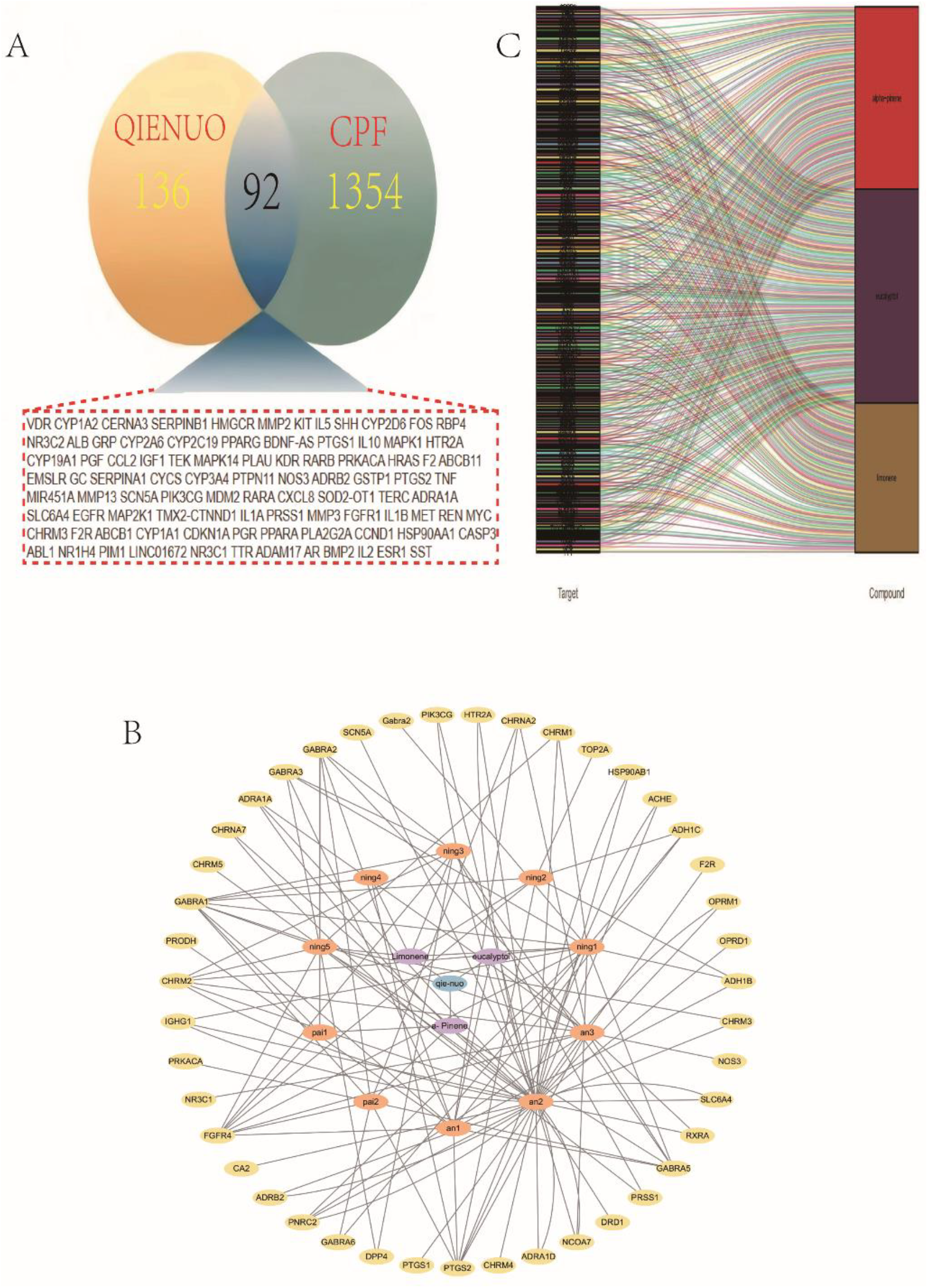
Identification of candidate targets through Protein-Protein Interaction (PPI) analysis. (A) PPI analysis. (B) PPI network based on cluster analysis using the MCODE plugin. (C) Identification of top 29 core targets based on DC ≥2 times the median DC. (D) Bar chart of the top 29 core targets.

### 3.3. Enrichment analysis of GO and KEGG

GO and KEGG enrichment of 92 “drug-disease” targets were analyzed using DAVID database. GO analysis was also carried out, and a total of 612 results were obtained, of which 477 were attributed to biological process (BP), 37 to cellular component (CC) and 98 to molecular function (MF). The top 10 items were displayed by creating circle diagram (Fig. 4A). The BP term with the highest enrichment degree is mainly related to external stimulus, negative regulation of cell population proliferation and positive regulation of cell population increment. The high concentration of CC is mainly concentrated in organelles, mitochondria and extracellular exosomes at the inner membrane boundary. GO analysis identified several biological processes related to the role of QIENUO in CPF, including positive regulation of transcription by RNA polymerase II, extracelluar exosome, chromatin binding and calmodulin binding. KEGG analysis showed that the targets of QIENUO -CPF were concentrated in the pathways closely related to QIENUO such as the Chimeric Virus, Cushing Syndrome and the related to melanin production. CystIn addition, highly enriched MF includes calmodulin binding, glucocorticoid receptor binding and chromatin binding. It is considered to be related to abnormal secretion of glucocorticoid and melanin. 155 signal pathways were identified by KEGG pathway enrichment analysis. Among the top 30 highly enriched KEGG pathways displayed by creating histogram and bubble chart (Fig. 4B), the highly enriched pathways related to hedgehog signal pathway, AMPK signal pathway, cGMP-PKG signal pathway, TGF-β signal pathway and cell cycle are closely related to the occurrence and progress of CPF.

**Fig. 4.**
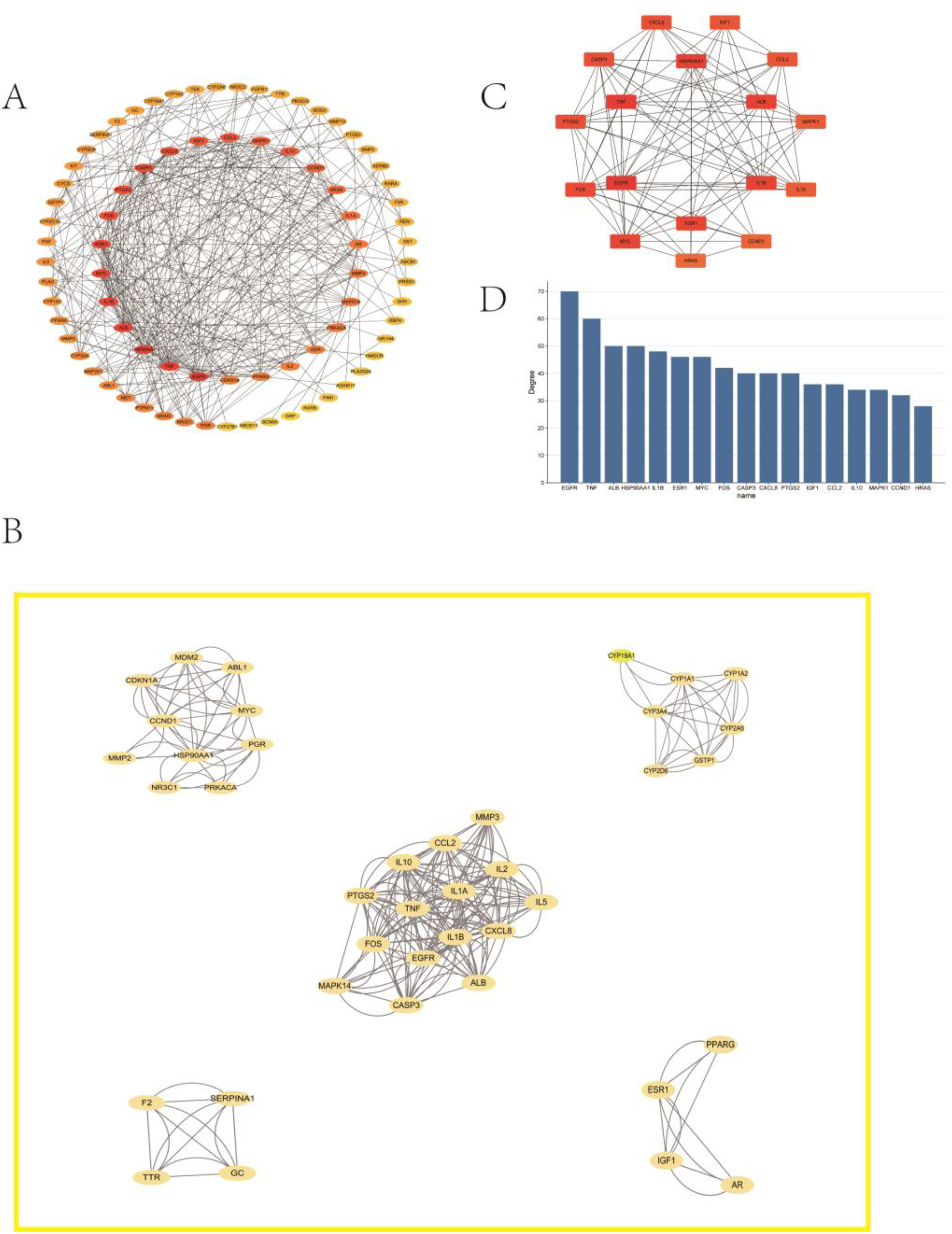
Results of GO and KEGG enrichment analysis (A) Barplot and bubble chart of GO functional enrichment analysis. (B) Barplot and bubble chart of the top 30 pathways based on KEGG enrichment analysis.

### 3.4. Compound-Target-Pathway Analysis and Identification of Core Targets of CPF and Molecular Docking

The drug-compound-target-pathway network was constructed with Cytoscape, and the top ten KEGG pathways were used to participate in the network construction. According to the network construction (Fig. 5A).The core genes IGF1, PTGS2, HSP90AA1, ESR1, MAPK1, CCL2, CCND1, IL1B, IL10, MYC, EGFR, ALB, CASP3, CXCL8, FOS, HRAS and TNF were screened. The core gene interaction network map was constructed by GeneMANIA database(Fig. 5B).The top five core proteins were screened out, namely ALB, EGFR, HSP90AA1, IL1B and TNF, which were only found in the components of eucalyptus oil. The three small molecular conformations of cineole are MOL005970, MOL000122 and MOL000917 respectively, so these five components are analyzed and docked with the three conformations of cineole.

**Fig. 5.**
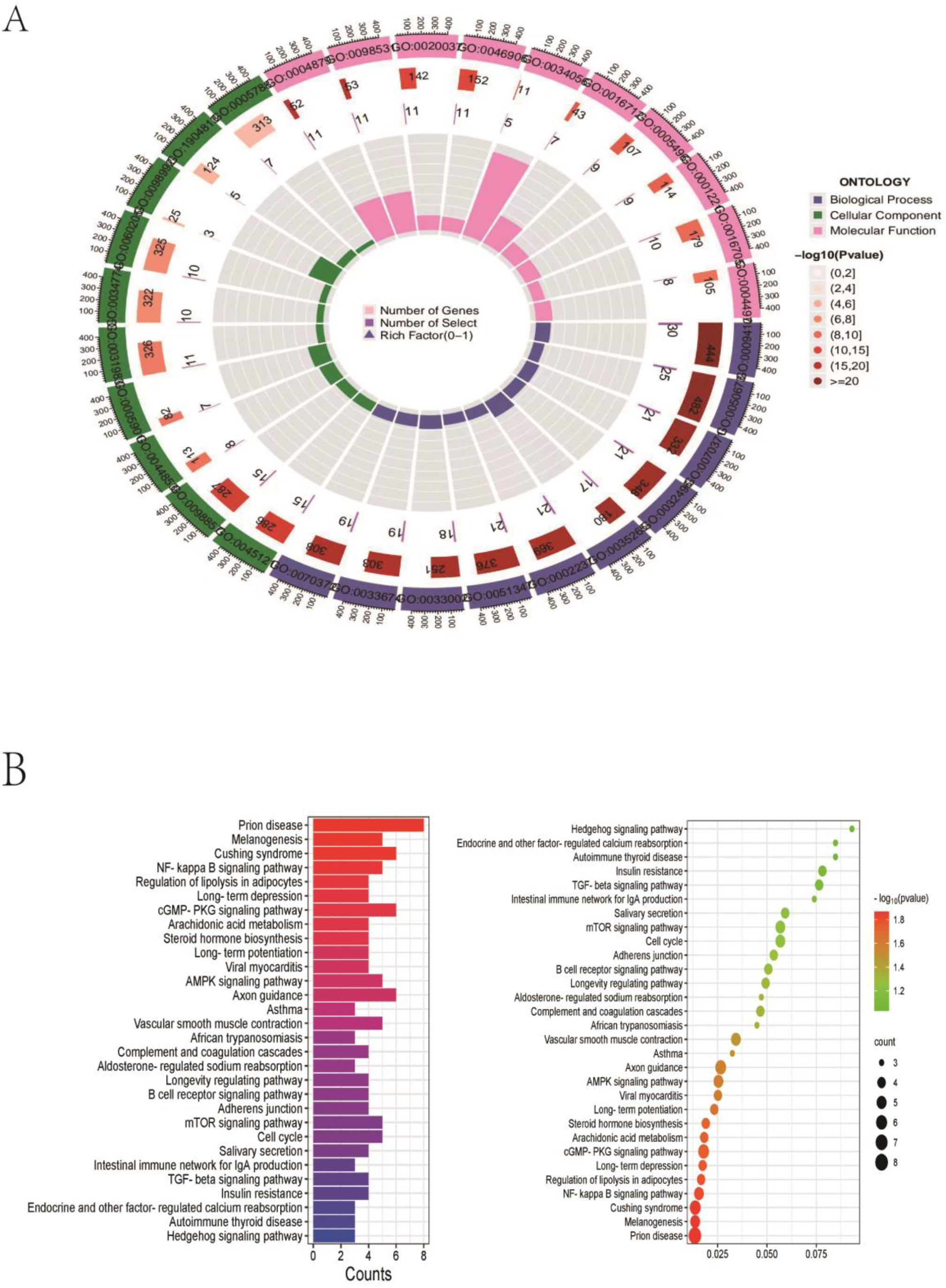
Construction of drug-compound-target-pathway network(A) Compound-target-pathway network.

**Fig. 6.**
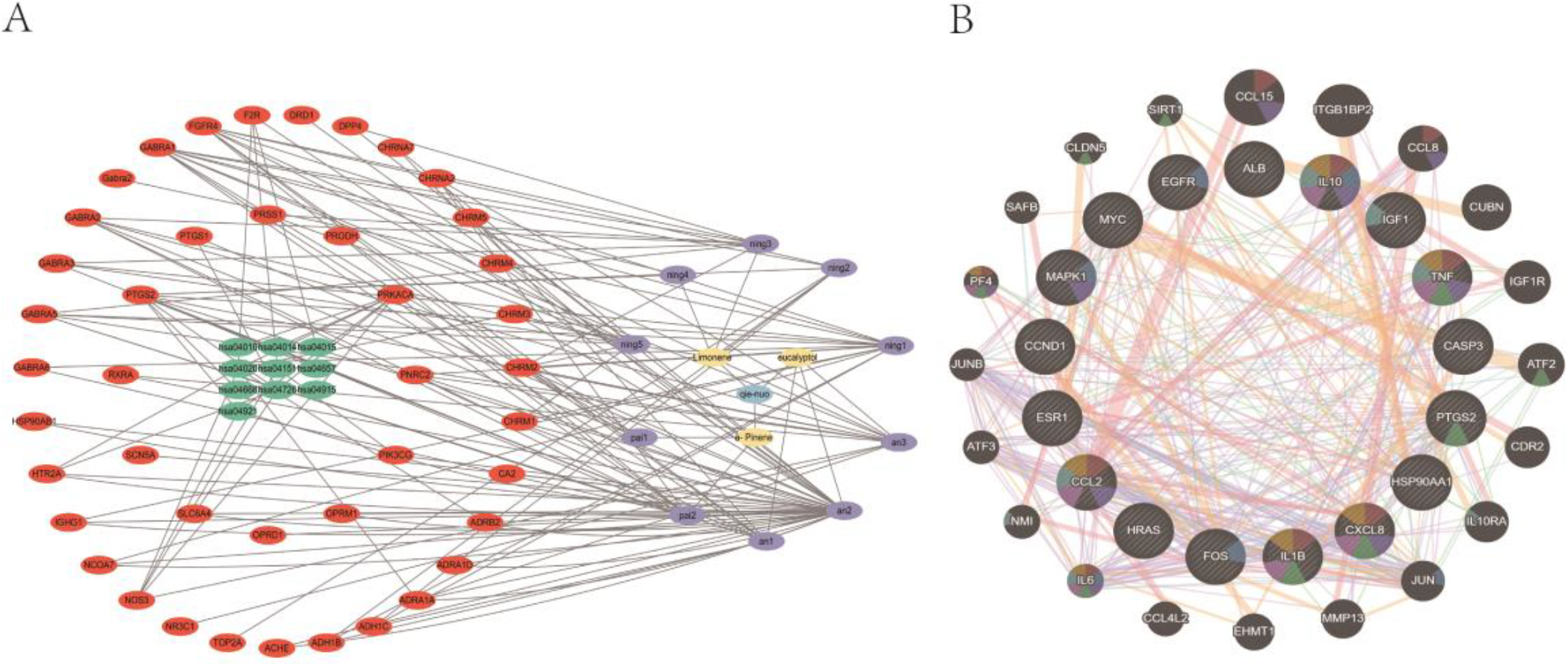
Docking patterns of core targets and active compounds of CPF

**Fig. 7.**
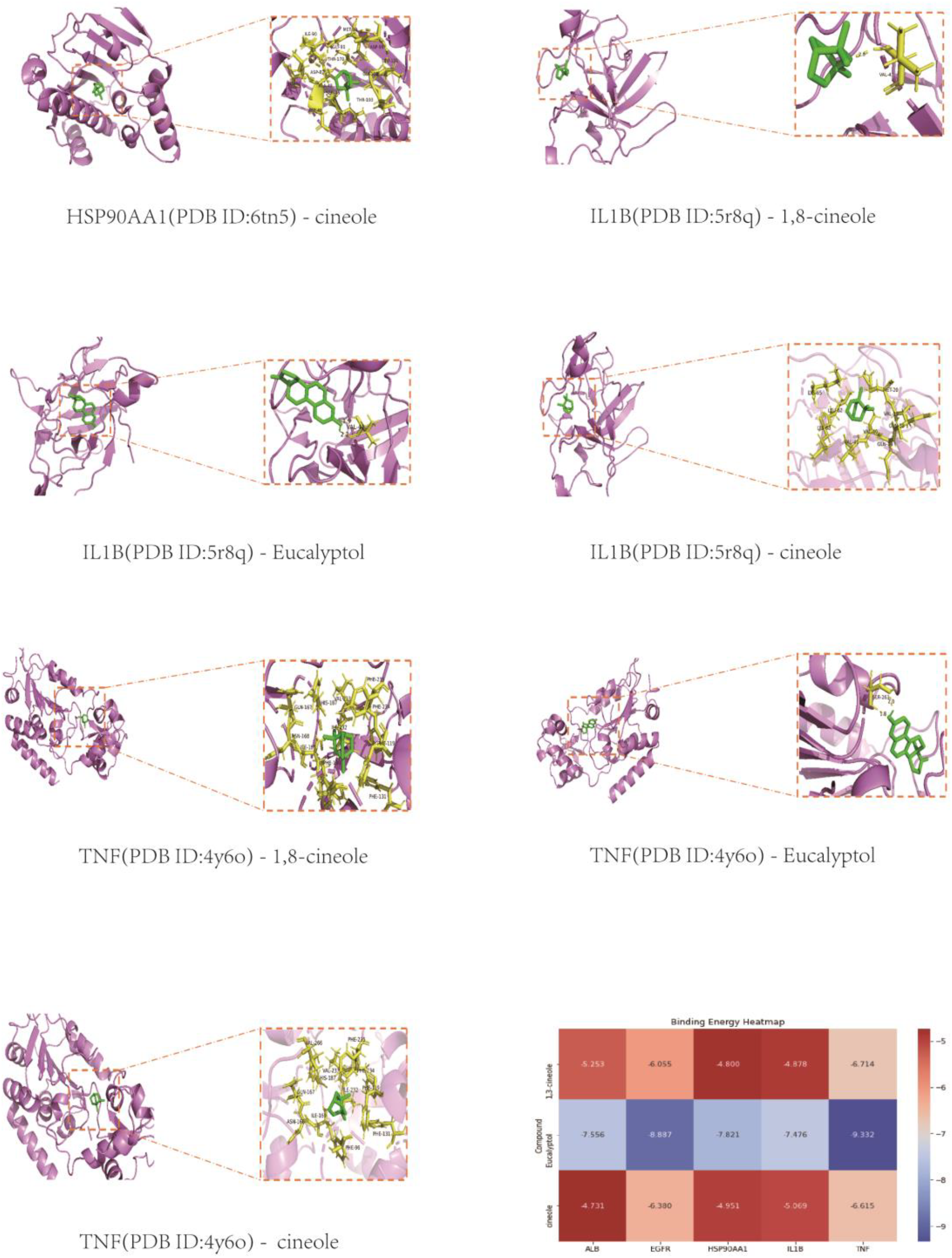
Other docking patterns of core targets and active compounds of CPF and Validation and screening of molecular docking. Heatmap of CPF molecular docking scores. Binding energies (kcal/mol) of core targets and active compounds of QIENUO.

### 3.5. Verification and visualization of molecular docking results

Molecular docking software AutoDockvina (https://vina.scripts.edu/) was used to study how the screened molecular compound ligands interact with the core target protein. The receptor protein used for molecular docking comes from the RCSB database. The results of thermogram show that when the binding energy of small molecules to protein is lower than -6kcal/mol, the target protein and compound have strong binding ability. There are 15 pairs of ligand-receptor pairs in the molecular docking of QIENUO -pulmonary cystic fibrosis, and the binding energy of 9 pairs of ligand-receptor pairs is lower than-6 kjmol-1. The binding energies are -7.556, -6.055, -8.887, -6.38, -7.821, -7.476, -6.714, -9.332 and -6.615, respectively (table 1). Among them, the binding energy of small molecule MOL005970 to protein is significantly lower than that of other small molecules, which indicates that small molecule MOL005970 is likely to be a potential binding target for QIENUO155in the treatment of CPF. In other related molecular docking results of QIENUO -CPF, the binding energy of MOL000917 to EGFR, MOL000122 and TNF is significantly higher than that of other target proteins to small molecules, which proves that there is also a strong interaction force, and the binding energy is -8.887kcal/mol and -9.332kcal/mol respectively. Some small molecules and hydrogen bonds are not formed or the binding energy is less than -6kcal/mol, which may be due to other forces between small molecules and adjacent amino acids or weak hydrogen bonds. All the results of molecular docking binding energy are presented in the form of thermogram. Finally, PyMOL is used to visualize the interaction and binding mode of compounds and targets with high free binding energy fraction. It can be seen from the figure that the interaction between small molecules and proteins is mainly hydrogen bonding, and the stability between the binding sites of small molecules and proteins is maintained by hydrogen bonding.

## 4. Discussion

Cystic fibrosis is the most common hereditary disease caused by gene defect of cystic fibrosis transmembrane conductance regulator. Pulmonary complications of chronic infection and eventual respiratory failure remain the most important threats. Until ten years ago, only symptomatic treatment was available^[19-21]^. However, since 2012, different combinations of cystic fibrosis transmembrane conductance regulator can be used for cystic fibrosis patients with different mutations. The appearance of these drugs has greatly improved the life expectancy and quality of life of patients with cystic fibrosis, but they still face great challenges in the possible complications in the later stage^[22]^. Among them, lung deterioration is a common event in children with cystic fibrosis^[23]^. The lung treatment of patients includes mucolytic agents (such as α-catenase), anti-inflammatory drugs (such as azithromycin) and antibiotics (such as tobramycin administered by atomizer)^[24,25]^, However, in clinical practice, the therapeutic effect of CPF is often unsatisfactory. Therefore, in clinical practice, it is very important to seek effective drugs to treat CPF and explore its potential mechanism.

As a compound preparation, Eucalyptol, Limonene and Pinene Enteric Capsules (QIENUO) are used as expectorants in clinic. It mainly contains eucalyptus oil essence, limonene and α -pinene Studies have proved that eucalyptus oil can alleviate cardiac fibrosis by inhibiting transient receptor potential anchor protein 1(TRPA1), and then inhibiting GRK5/NFAT signal transduction^[26]^. Animal experiments have also proved that Chenopodium may have a strong therapeutic effect on CPF but its therapeutic mechanism is still unclear. Therefore, it is necessary to analyze the interaction mechanism between QIENUO and CPF through bioinformatics and molecular docking.

It is worth noting that the target of QIENUO related to CPF is display from the compound-target-pathway network, and then they are verified by molecular docking. This will help to more clearly understand the mechanism of how QIENUO treats CPF. Specifically, our study used molecular docking method to evaluate the interaction between five key target proteins related to CPF (EGFR, TNF, ALB, HSP90AA1, IL1B) and three active molecular compounds (eucalyptol, Limonene, a-Pinene). According recent research, two key proteins play an important role in the treatment of CPF. Chronic Obstructive Pulmonary Disease(COPD) and CPF are interrelated through the EGFR-ADAM17 pathway^[27]^,maybe CPF occurs with COPD. Another research prove that a role for genes of the TNF-alphaR/NFkappaB pathway in controlling the proinflammatory state of cystic fibrosis lung epithelium^[28]^.

Five core proteins are all from eucalyptus oil, the binding affinity of EGFR, TNF, ALB, HSP90AA1, IL1B and eucalyptus is obviously higher than that of Limonene and a-Pinene, which can be well explained by the results of molecular docking^[29]^. It has been previously reported that cineole can alleviate bleomycin-induced pulmonary fibrosis by regulating the polarization of M2 macrophages, which in turn inhibits the activation of signal molecules (for example, STAT6 and p38MAPK) and the expression of transcription factors (for example, KLF4 and PPAR-γ) ^[30-33]^. Therefore, eucalyptol may be a potential therapeutic agent for CPF^[34]^. Our research found that eucalyptol formed a stable hydrogen bond structure with almost all small molecular ligands. As a result of molecular docking^[35]^, TRP-147, ARG-97, ASP-855, MET-799, ARG-841, LEU-101, HIS-148, ASP-48, VAL-41 are the site where hydrogen bonds are formed on amino acids, could be a key point for new drug research. Further study on the interaction between eucalyptol and these proteins may help to develop targeted therapy for CPF.

## 5. Conclusion

Our research takes the lead in using bioinformatics methods, including network pharmacology and molecular docking, to study the pharmacology and molecular mechanism of QIENUO in treating CPF. The evidence obtained by bioinformatics and computer analysis shows that eucalyptus oil may be the main bioactive component in the treatment of CPF with QIENUO. In addition, the therapeutic effect of QIENUO on CPF includes many pathways, such as Positive regulation of Transcription by RNA polymerase II, extracelluar exosome and chromatin binding. In a word, this study analyzed the multi-component and multi-pathway characteristics of QIENUO, revealed the potential mechanism of QIENUO, and provided valuable resources for the application and research of QIENUO in CPF.

## Conflict of interest statement

The authors declare no competing interests.

## Author contributions

Y.Z. conceived and designed the current study. T.T., X.Z., and W.C. performed analyses. T.T. and Y.Z. prepared the manuscript. C.W and Y.Z. reviewed and edited the manuscript. All authors read and approved the manuscript.

## Funding information

Y.Z. was supported by Discipline Cluster of Oncology, Wenzhou Medical University, China (No.z2-2023024).

## Figure legend

Table 1. This table including all building energy, unit: kcal/mol

## Notes

### Competing Interest Statement

The authors have declared no competing interest.

