## Supplementary material for "Analyzing the Molecular Mechanism of Eucalyptol, Limonene and Pinene Enteric Capsules (QIENUO) in the Treatment of Pulmonary cystic fibrosis with Network Pharmacology and Verifying Molecular Docking": table1

|  | Receptor | ALB | EGFR | HSP90AA1 | IL1B | TNF |
| --- | --- | --- | --- | --- | --- | --- |
| Ligand  1,3-cineole  Eucalyptol  cineole |  | -5.253  -7.556  -4.731 | -6.055  -8.887  -6.38 | -4.8  -7.821  -4.951 | -4.878  -7.476  -5.069 | -6.714  -9.332  -6.615 |

Table 1. This table including all building energy , unit: kcal/mol
